# Multifactorial remodeling of the cancer immunopeptidome by interferon gamma

**DOI:** 10.1101/2022.03.23.485466

**Authors:** Alice Newey, Lu Yu, Louise J Barber, Jyoti S. Choudhary, Michal Bassani-Sternberg, Marco Gerlinger

## Abstract

IFNγ alters the immunopeptidome presented on HLA class I (HLA-I), and its activity on cancer cells is known to be important for effective immunotherapy responses. We performed proteomic analyses of untreated and IFNγ-treated colorectal cancer patient-derived organoids (PDOs) and combined this with transcriptomic and HLA-I immunopeptidomics data to dissect mechanisms that lead to remodeling of the immunopeptidome through IFNγ. IFNγ-induced changes in the abundance of source proteins, switching from the constitutive- to the immunoproteasome, and differential upregulation of different HLA alleles explained some, but not all, observed peptide abundance changes. By selecting for peptides which increased or decreased the most in abundance, but originated from proteins with limited abundance changes, we discovered that the amino acid composition of presented peptides also influences whether a peptide is up- or downregulated on HLA-I through IFNγ. The presence of proline within the peptide core was most strongly associated with peptide downregulation. This was validated in an independent dataset. Proline substitution in relevant core positions did not influence the predicted HLA-I binding affinity or stability, indicating that proline effects on peptide processing may be most relevant. Understanding the multiple factors that influence the abundance of peptides presented on HLA-I in the absence or presence of IFNγ is important to identify the best targets for antigen-specific cancer immunotherapies such as vaccines or T-cell receptor engineered therapeutics.

## Introduction

The presentation of peptides on HLA class I (HLA-I) is central for the adaptive immune system to detect malignant cells. Presentation of immunogenic peptides such as non-mutated cancer-associated antigens or neoantigens on malignant cells facilitates their recognition and destruction by cytotoxic CD8 T-cells. Interferon gamma (IFNγ) is a cytokine that is released from activated CD8 T-cells and other immune cell types. IFNγ binds to IFNγ receptors, which activate the JAK/STAT pathway, leading to expression of interferon response factor transcription factors (IRFs). IRFs stimulate the expression of a plethora of IFNγ-regulated genes leading to major changes in the cellular transcriptome and proteome. Proteins involved in the processing and subsequent presentation of peptide antigens on HLA-I molecules show particularly strong upregulation, including the immunoproteasome catalytic components PSMB8, PSMB9, and PSMB10 which facilitate an increase in overall proteasomal activity, and also a specific increase of its chymotryptic activity(1,2). Peptidases which can trim, but also destroy, peptides before loading onto HLAs, such as LAP3(3), THOP1(4), ERAP1(5), and ERAP2(6), and the peptide transporters TAP1 and TAP2, which shuttle peptides into the endoplasmic reticulum where HLA loading occurs, are also upregulated. Furthermore, IFNγ increases HLA expression. The combined effect of increased proteasomal peptide generation, peptide processing and transport, and HLA upregulation is a strong increase of peptide presentation by HLA-I on the cell surface. Further to this, IFNγ exposure inhibits the cell cycle and triggers apoptosis(7).

In contrast to these anti-tumour effects, IFNγ also promotes the expression of immunosuppressive molecules. These include PD-L1, the ligand of the PD1 immune checkpoint, and IDO1, whose expression in cancer cells and other cells in the tumour microenvironment suppresses T-cell activity(8–10). Immunotherapy with PD1/PD-L1 inhibitors has been highly successful in several cancer types(11–13). This supports a dominant role of the PD1/PD-L1 immune checkpoint in restraining tumour reactive T-cells. Consistent with a central role of IFNγ for PD-L1 expression, tumours that respond to PD1/PD-L1 inhibitors often show high IFNγ activity(14). Moreover, several recent studies have shown that defective IFNγ-signaling in cancer cells leads to resistance to immunotherapy with checkpoint-inhibitors(15–17).

Although intact IFNγ signaling in cancer cells is critical for checkpoint-inhibitor efficacy, it is still unclear which specific IFNγ-induced molecular changes are responsible for this dependency. Understanding how the immunopeptidome is remodeled by IFNγ in greater detail may provide insights into this. Furthermore, novel immunotherapies such as cancer vaccines(18,19) and engineered TCR-based therapies such as Tebentafusp(20), target T-cells towards specific peptide antigens presented on HLA of cancer cells. Understanding the characteristics of antigens that are consistently presented in the presence or absence of IFNγ, and which ones are lost or sparsely presented in one of these conditions, hence appears highly relevant for the selection of optimal target antigens.

Previously, we studied the immunopeptidome of 5 colorectal cancer (CRC) patient-derived organoids (PDOs) by mass spectrometry (MS). PDO cells were grown to large numbers followed by immunoaffinity capture of HLA-I/peptide complexes, analysis by high performance liquid chromatography and tandem MS. This detected between 2,124 and 16,030 HLA-I peptides per PDO(21). Treatment of PDOs with IFNγ strongly increased HLA-I expression (mean 219.5% increase) but only had a modest effect on the number of unique peptides presented (mean 7.1% increase). However, a much larger number of peptides changed in abundance, and between 1,439 and 3,942 peptides were gained, and 561 to 2,446 peptides were lost on individual PDOs through IFNγ.

Furthermore, we found that peptides generated by chymotryptic-type cleavage activity were more likely to increase in abundance, which we attributed to the switch of the proteasome to immunoproteasome triggered by IFNγ signaling. However, we did not detect a strong enrichment of chymotryptic-like ligands in the peptides which most-highly increased in abundance under IFNγ. This highlights a limited understanding of the molecular mechanisms and peptide features that regulate peptide abundance on HLA-I in IFNγ conditions. The aim of this work was to dissect the mechanisms that lead to up-/downregulation or appearance/loss of specific peptides under IFNγ exposure. We combined global cellular proteomic analysis with our published transcriptomics and immunopeptidomics datasets(21) to first investigate the impact of transcript and protein abundance on immunopeptidome remodeling, and to subsequently analyze peptide regulatory mechanisms that are independent of source protein abundance. The insights from this study should ultimately lead to more accurate predictions of the immunopeptidome in cells exposed to IFNγ, information which could be valuable for cancer vaccine or TCR therapy designs.

## Materials and methods

### PDO culture and treatment

Established PDOs were expanded to large numbers (3.85×10^7^–1×10^8^ cells/pellet) in DMEM/F12 media with 20% fetal bovine serum, 1X Glutamax, 100 units/ml penicillin/streptomycin and 2% matrigel. For treatment, cells were changed into fresh media supplemented with DMSO or 600ng/mL IFNγ (R&D Systems) and incubated for 48 hours. Cells were harvested with TrypLE express (ThermoFisher). PDOs were cultured identically for transcriptomic, proteomic, and flow cytometric analysis.

### RNA-sequencing

We reanalyzed our previously described RNA sequencing data(21,22).

### Tandem-mass-tag (TMT) proteomics

PDOs were cultured as described, washed twice with ice-cold PBS and snap-frozen before further processing. Further technical details are provided in the Supplementary Methods. The protein abundance values in each sample were normalized by the loading input of each sample.

### MS immunopeptidomics

MS immunopeptidomics data had been acquired as described(21,23). The peptide intensity values in each sample were normalized by the total detected peptide intensity for each condition. The validation dataset was obtained from(24) and was normalized in the same way.

### Prediction of NetMHC percentile ranks from MS-detected peptides

All HLA-I MS-detected peptides were entered into NetMHCpan4.0(25). The HLA allotypes determined for each PDO line were used to subset peptides in to HLA-I-allocated groups. nM prediction was used where peptide HLA-affinity was the factor in question, and predicted rank was used where all peptide presentation factors were to be considered.

### Relative peptide start position

Relative peptide start position within protein was calculated for each peptide by dividing the peptide start position by the protein length. The longest protein length for each protein was selected from the database search Fasta file to ensure every peptide was encompassed.

### HLA typing

HLA typing results from the previous publication were used(21).

### Surface HLA quantification by flow cytometry

HLA surface expression was assessed using pan-HLA-A/B/C antibody (BioLegend, W6/32), anti-HLA-A03 (ThermoFisher Scientific, GAP.A3), and anti-HLA-B27 (Bio-Rad, HLA-ABC-m3). Samples were run on a Sony SH800 cell sorter.

### Statistics

Statistical calculations and plots were performed in R (www.r-project.org) and on GraphPad Prism v9. Z-scores for the amino acid enrichment analysis were calculated by subtracting each value by the mean of all the difference values, then dividing by the standard deviation of all the difference values.

## Results

The aim of this study was to elucidate the molecular mechanisms through which IFNγ alters HLA-I peptide presentation by comparing the immunopeptidome of untreated and 48-hour IFNγ-treated CRC PDOs (CRC-01, CRC-04, CRC-05). This was achieved by combining previously generated transcriptomics and immunopeptidomics data(21) with new global proteomics data, obtained by tandem-mass-tag MS (TMT-MS). 7,408 proteins were detected by TMT-MS across the three PDOs. IFNγ-induced fold-changes (FC) of transcript and protein abundance (Figure 1A) showed a significant positive correlation (Spearman’s r=0.34-0.69, p<2.2×10^−16^ for all 3 PDOs). IFNγ increased the expression of a large number of genes/proteins whereas downregulation was only apparent in a smaller number of genes/proteins and was of limited magnitude.

**Figure 1:**
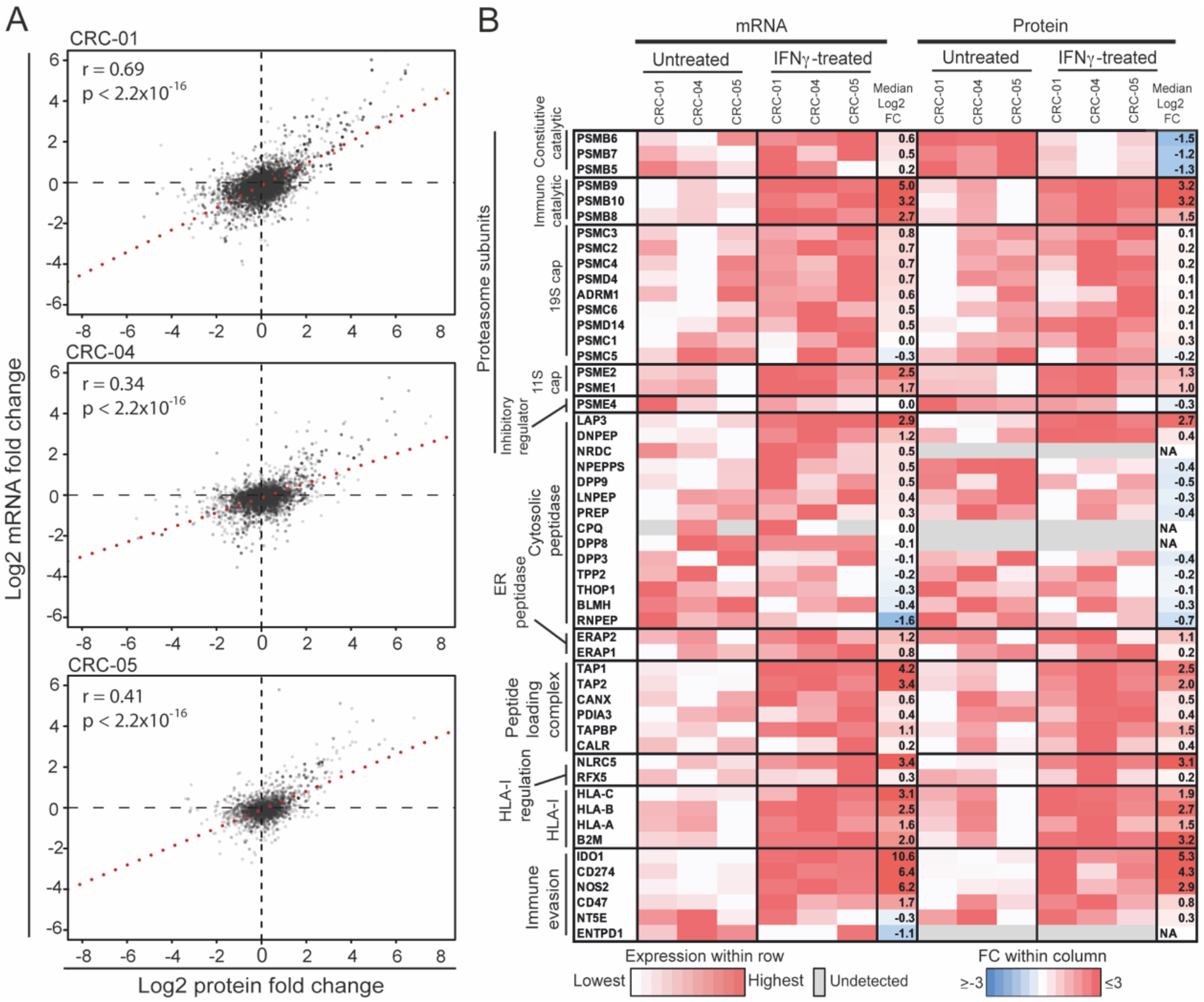
Transcriptomic and proteomic changes with IFNγ treatment. **A:** Correlation of the fold-change in normalized mRNA read numbers against the fold-change in normalized protein intensity. The Spearman’s rank test was used for statistical analysis. **B:** mRNA expression and protein intensity of selected genes in untreated and IFNγ conditions.

We next assessed whether transcripts/proteins that were previously described as IFNγ-regulated, *and* have roles in antigen processing and presentation or immune evasion, undergo the expected changes. IFNγ treatment increased RNA expression of most proteasome components, including constitutive proteasome catalytic subunits (PSMB5-7), and immunoproteasome catalytic components (PSMB8-10) (Figure 1B). In contrast, proteome data showed a strong decrease of the constitutive catalytic subunits. This disparity between RNA and protein abundance can be explained by the fact that immunoproteasome assembly is four times faster than that of the constitutive proteasome(26), so the excess unbound constitutive catalytic components will be degraded(1). This switch from constitutive to immunoproteasome alters the cleavage specificity towards an increased chymotryptic activity, as we observed in our previous study(21). The regulatory caps of the proteasome also change with IFNγ treatment; in the absence of IFNγ, the 26S proteasome forms by addition of the 19S cap to each end of the 20S proteasome core. The 19S cap is responsible for binding polyubiquitinated proteins and actively transporting them in to the 20S proteasome core. IFNγ increased the expression of the 11S cap subunits which facilitate ubiquitin-independent proteasomal degradation of proteins and enhance proteasomal throughput (27–29). PSME4 is a another proteasome cap subunit, recently shown to inhibit the production of HLA-I compatible peptides(30). The decrease in PSME4 protein abundance through IFNγ may further increase peptide production.

Among cytosolic peptidases, LAP3 (with a cleavage specificity towards hydrophobic N-terminal amino acids, primarily leucine) increased in protein abundance (Figure 1B), and most other cytosolic peptidases showed a small decrease. Both endoplasmic reticulum N-terminal aminopeptidases ERAP1 and ERAP2, which help to shape the immunopeptidome by final trimming on HLA-I(5,6), increased. TAP transporters and peptide loading complex (PLC) components increased in RNA expression and protein abundance. Further to this, NLCR5, the master transcription factor for HLA-I expression, and consequently HLA-A, -B and -C increased strongly.

We also assessed whether genes and proteins that are known to inhibit the activity of immune cells were upregulated by IFNγ. Most immune evasion genes increased in expression, with IDO1 and CD274/PD-L1 showing the strongest increase at the protein level. Thus, IFNγ triggered expected changes in known IFNγ-regulated genes across all three organoid lines.

We next assessed to what extent a change in abundance of source proteins influenced the presentation of their derived peptides on HLA-I. We plotted the FC in protein abundance between untreated and IFNγ conditions against normalized FC of all HLA-I presented peptides (Figure 2A). Normalization of the immunopeptidome data was performed similar to the normalization approach used for RNA and protein expression data: peptide intensity values were normalized so that the total intensity in untreated and IFNγ treated conditions were identical for each PDO. This allowed us to investigate peptide abundance changes beyond those driven by the absolute increase in HLA-I expression with IFNγ. This analysis revealed two distinct components in each PDO: one group of up- or downregulated peptides showed concordant changes in the source protein abundance through IFNγ, and a second group of up- or downregulated peptides derived from proteins with no or limited change in abundance (log2 -1 to 1 FC). This shows that source protein abundance is one important driver of peptidome remodeling, but also that the surface presentation of an even larger number of peptides is controlled by additional mechanisms.

**Figure 2.**
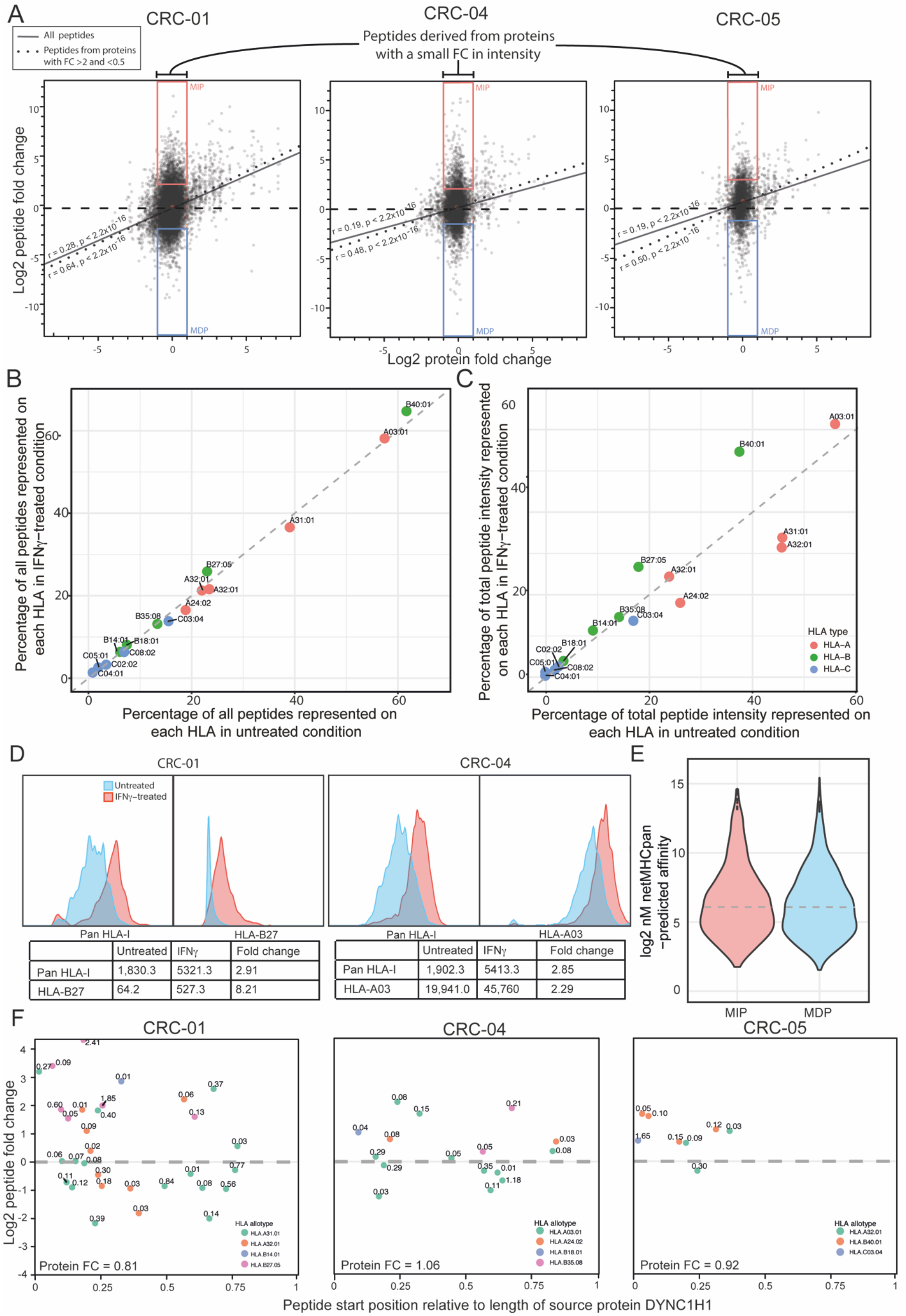
Influence of protein abundance and HLA expression changes on immunopeptidome remodeling. **A:** Correlation of protein fold-change between untreated and IFNγ conditions, against normalized immunopeptidomics fold-change. Regression lines for all peptides are displayed as a solid black line, regression lines for peptides from proteins with FC <0.5 or >2 as a dotted line. The Spearman rank test was used for significance testing. Most increasing peptides (MIPs - top 10^th^ percentile peptide FC) and most decreasing peptides (MDPs - bottom 10^th^ percentile peptide FC) derived from low fold-change proteins (0.5-2X fold-change) are highlighted with red and blue boxes, respectively. **B:** Percentage of all peptides per PDO that were attributed to each HLA by NetMHCpan4.0 in untreated vs. IFNγ-treated conditions. **C:** Percentage of the total peptide intensity per PDO represented on each HLA by NetMHCpan4.0 in untreated vs. IFNγ-treated conditions. **D:** Expression of total surface HLA-I and single HLA-I allotypes in organoid lines CRC-01 (HLA-B27) and CRC-04CRC-04 (HLA-A03), measured by flow cytometry. **E:** Log2 NetMHCpan4.0-predicted nM binding affinity values for MIPs vs MDPs. The median is marked with a dotted line. **F:** Log2 change in peptide intensity between untreated and IFNγ-treated conditions for peptides derived from the protein DYNC1H1, plotted against the relative position of the peptide in the protein. Peptides are color-coded by their NetMHCpan4.0-predicted source HLA, with the NetMHCpan4.0 predicted rank annotated above. The fold change of the DYNC1H1 protein in each PDO is noted at the bottom.

Our next aim was to understand how this second group of peptides is regulated in these PDOs. We first hypothesized that IFNγ upregulates different HLA alleles by different levels and that this may affect the diversity or abundance of their corresponding peptide repertoires. Plotting the number of unique peptides that were predicted by NetMHCpan4.0(25) to bind each HLA allele, showed only a small changes through IFNγ treatment (Figure 2B). Next, we plotted the total peptide intensity per HLA allele as a surrogate measure of the total number of peptides presented on each HLA. When treated with IFNγ, the relative intensity of peptides presented on HLA-B allotypes increased, whereas those on 3 of 5 HLA-A allotypes decreased (Figure 2C). This indicates that IFNγ treatment increased the percentage of peptides presented by HLA-B among all HLA-I presented peptides, and reduced that presented by HLA-A. Importantly, the absolute number of peptides presented on HLA-A may still increase as HLA-A, -B and –C all showed upregulation in the proteomic data. To validate these findings, we performed flow cytometry staining for total HLA-I and for two HLA allotypes (HLA-A03 and HLA-B27), for which specific monoclonal antibodies were available. Pan-HLA antibody staining on CRC-01 showed a 2.91-fold upregulation of total HLA, whereas staining for HLA-B27 showed an 8.21-fold increase (Figure 2D), 2.82-fold more than the total HLA. Further to this, CRC-04 showed a 2.85-fold upregulation of total HLA and 2.29-fold upregulation of HLA-A03, which is 0.8-fold that of the total HLA upregulation (Figure 2D). This validated the results from the peptide analysis and is consistent with previous studies showing a stronger upregulation of HLA-B compared to HLA-A molecules with IFNγ(23,31,32). Thus, peptides on HLA-B allotypes increase in relative abundance within the immunopeptidome of individual PDOs, whereas those on HLA-A showed a relative decrease with IFNγ treatment.

We next assessed whether binding affinities of peptides to their cognate HLA allele may influence up- or downregulation. We focused on 9-mer peptides that changed strongly in intensity, yet originated from source proteins with modest abundance changes (defined again as log2 -1 to 1 FC). The group of Most Increasing Peptides (MIPs) was defined as those in the top 10^th^ percentile of the immunopeptidomics FC data (red box in Figure 2A) and Most Decreasing Peptides (MDPs) as the bottom 10^th^ percentile (blue box in Figure 2A). Plotting the NetMHCpan4.0-predicted affinities of MIPs and MDPs for their HLA (Figure 2E) revealed a similar data distribution between the two groups, and no significant difference between medians (p=0.1204, Mann-Whitney test). Thus, peptide binding affinity did not noticeably impact whether a peptide was up- or downregulated by IFNγ.

Our next hypothesis was that other peptide-specific factors determine up- or down-regulation, independently of protein and HLA abundance changes. To assess this, we first focused on large proteins that each contributed multiple MS-detected peptides in individual PDOs. A single protein can undergo a specific change in abundance and turnover with IFNγ treatment and this should affect all peptides that originate from that protein similarly. We therefore reasoned that the detection of up- and downregulation of peptides from the same protein with IFNγ treatment would indicate that peptide-specific characteristics influence these abundance changes. One limitation of this approach is that it does not control for differences in protein isoforms, which may be relevant for some proteins. Analysis of DYNC1H1, the protein that contributed the largest number of peptides across each of our PDOs, (Figure 2F), showed that some peptides originating from the same protein increased, whereas others decreased with IFNγ treatment. This appeared independent of the cognate HLA allotype, and is hence not the consequence of differential HLA-A and –B upregulation. Analysis of 9 additional large proteins, showed similar results (Supplementary Figure 1A-J). Thus, peptide-specific factors play a major role in determining whether a peptide is up or downregulated through IFNγ.

Some publications identified an overrepresentation of peptides derived from the N-terminus of a protein due to premature termination of translation or nonsense mediated decay, but whether this effect increases or decreases with IFNγ treatment is unknown(33,34). No systematic increase or decrease in the abundance of peptides located closer to the N-terminus was apparent in the analysis of long proteins (Figure 2F, Supplementary Figure 1A-J). To assess a much larger number of data points, we plotted the frequency of MIPs and MDPs against their relative location in the source protein (Supplementary Figure 1I). This showed only modest differences, suggesting that location within the protein had little effect on peptide production between UT and IFNγ conditions.

We next investigated whether the amino acid composition of MIPs or MDPs influenced IFNγ-induced changes in peptide abundance. MIPs and MDPs from each PDO were subset into their NetMHCpan4.0-predicted source HLA. We then analysed the frequency of all amino acids at each position of the 9-mers and values for MDPs were subtracted from MIP values (Figure 3A). The amino acid composition of the N-terminal and the C-terminal extensions adjacent to the presented peptide may also influence peptide processing(35), for example through the presence of specific cleavable amino acids or motifs for proteasome or peptidase processing. Therefore, we also assessed the 9aa N- and C-terminal extensions. Most differences were small, but Z-score analysis identified 12 amino acids in specific positions where the difference was 2.5 times larger than the standard deviation of all values (dotted outlines in Figure 3A).

**Figure 3.**
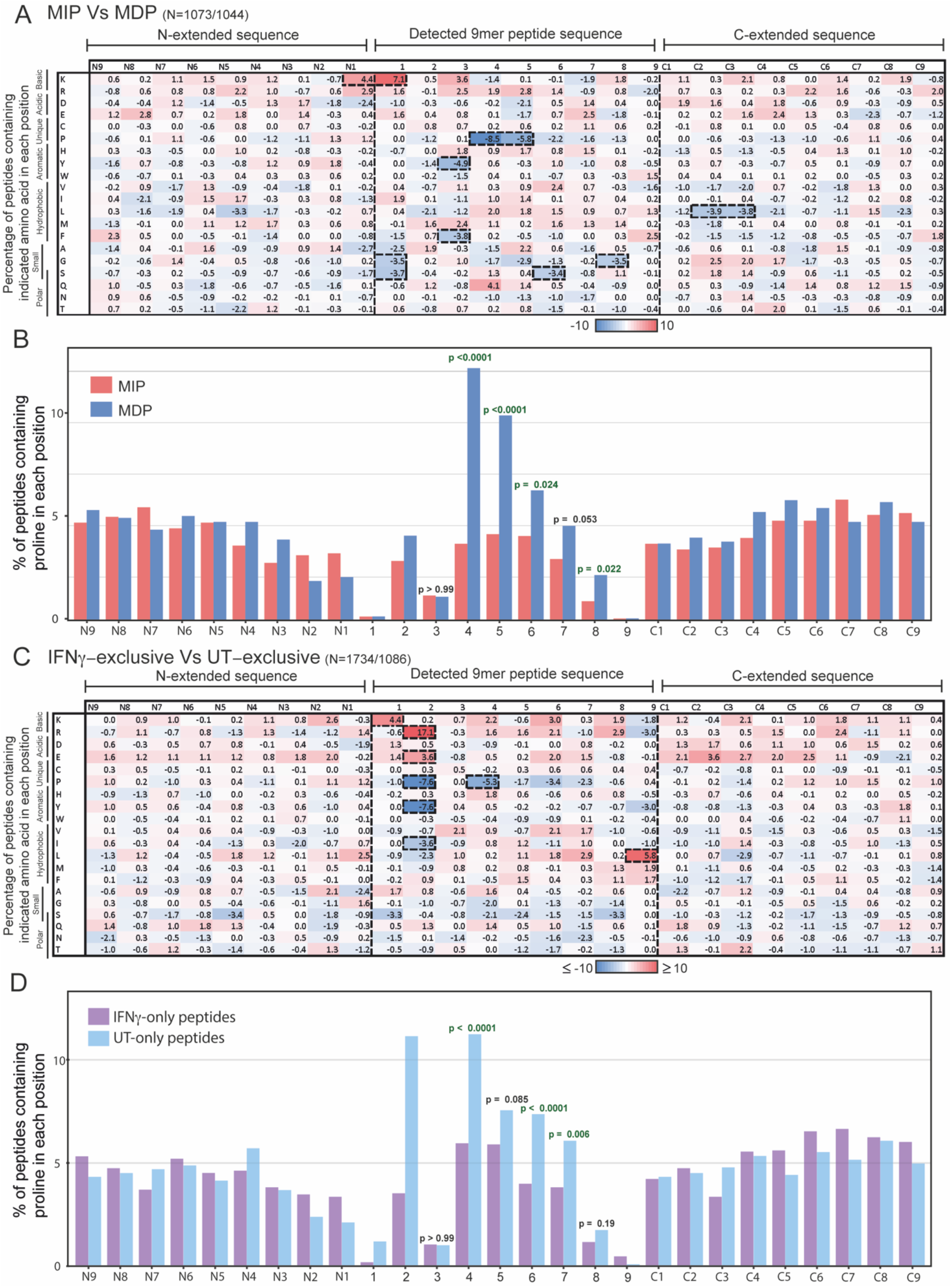
Amino acid composition of MIPs vs MDPs and UEPs vs IEPs. **A:** Heatmap of the amino acid composition changes between 9-mer Most increasing peptides (MIPs - top 10^th^ percentile peptide FC) and (MDPs - bottom 10^th^ percentile peptide FC) as well as N- and C extensions. Detailed peptide numbers provided in Supplementary Table 1 and 2. Percentage of peptides with highlighted amino acid in each position were calculated for each group, then the percentage values for the MDPs (N= 1,073) were subtracted from the MIPs (N=1,045). **B:** Percentage of peptides with proline in each position for MIPs (N= 1,073) and MDPs (N=1,045). **C:** Heatmap of the amino acid composition changes between (IFNγ-exclusive peptides – IEPs) and (untreated-exclusive peptides – UEPs). Percentage of peptides with each amino acid in highlighted position were calculated for each group, then the percentage values for the UEPs (N=1,195) were subtracted from the IEPs (N=1,909). **D:** Percentage of peptides with proline in each position in IEPs and UEPs. Z-score analysis was used for the heatmaps, and changes with a Z-score <-2.5 or >2.5 were highlighted with a thick dotted line. Fisher’s exact test was used for statistical analysis of the proline abundance values, with significant results are indicated in green.

There was an overrepresentation of lysine, a basic amino acid, in position N1 of the N-terminal extension, with a Z-score of 3.4. Notable in this context was also the less marked overrepresentation of arginine, another basic amino acid, in position N1 which fell below the Z-score cutoff, with a score of 2.3. Although the increase in basic residues in position N1 may suggest an increase in tryptic cleavage activity with IFNγ, we also identified an overrepresentation of lysine (Z-score of 5.5) in position 1 of MIPs. This cannot be explained by tryptic activity, as tryptic-like activity cleaves to the C-terminus of a basic residue, not the N-terminus. The small amino acids glycine and serine were underrepresented in position 1 of the MIPs (Z-scores of -2.7, and -2.9, respectively). This may be caused by the exchange of PSMB5, whose binding pocket has a preference for small amino acids, for PSMB8 in the immunoproteasome, which has a preference for chymotryptic-like substrates(2). However, some of these changes may also be due to activity of peptidases, as peptidases commonly cleave away additional amino acids from the N-terminus of a longer precursor peptide(35).

The only positions in the C-terminal extension highlighted by our Z-score analysis were positions C2-C3, in which leucine was underrepresented (with Z-scores -3 and -2.9). The widely accepted view is that the C-terminus of the peptide is directly generated by the proteasome(23,36), as no carboxypeptidases have been identified in antigen processing. Therefore, one possible explanation for our observations is that the leucines in C2 and C3 influence pepide generation by the immunoproteasome.

The amino acid with the largest difference within the 9-mer peptides was proline. This was underrepresented in positions 4 and 5 of MIPs with a Z-scores of -6.6 and -4.5, respectively (Figure 3A). Underrepresentation continued in the consecutive positions 6-8, but these did not cross the Z-score threshold (−1.7, -1.2, and -1.0, respectively). We next plotted the proline abundance in MIP and MDPs for each position of the peptide and its N- and C extension and applied statistical testing. This showed significant differences for positions 4-6 and 8 (Figure 3B). We furthermore analysed all 10-mer MIP and MDP peptides identically to the 9-mers to ascertain whether these observations could be reproduced. This confirmed a similar underrepresentation in MIPs in positions 4-7, crossing the Z-score threshold in positions 4 and 5 (Supplemental Figure 2A). To assess whether the depletion of proline in MIPs was detectable across PDOs and different HLA allotypes, we furthermore analysed the proline abundance in each position of 9-mer peptides separately for each PDO and HLA allotype. This showed consistent proline underrepresentation in MIPs, most strongly in positions 4 and 5 (Supplemental Figure 2B). Thus, peptides with proline in positions 4-5 were more likely to be downregulated through IFNγ treatment and this was neither a PDO, nor a HLA-specific effect. Prolines in position 6-8 appeared to have a similar, but less pronounced effect. The heatmaps further demonstrated that as proline decreased in abundance, there was no corresponding increase in another amino acid, but small increases dispersed among several amino acids. This suggests that the decrease of proline is a specific effect of IFNγ.

We also detected other larger changes in Z-score within the peptide: -4.9 and -3.8 for tyrosine and phenylalanine in P3; -3.4 for serine in P6; and -3.5 for glycine in P8. However, with the exception of tyrosine in P3, these findings were not reproduced in the original positions, or original positions +/-1 in the analysis of 10-mers (Supplementary Figure 2A).

We next assessed whether peptides that were only detected in untreated PDOs (untreated-exclusive peptides – UEPs) or only detected with IFNγ treatment (IFNγ-exclusive peptides – IEPs), and derived from proteins with a low FC (log2 -1 to 1 FC) showed the same signal. Proline was again underrepresented at positions 4-7 in IEPs compared against the UEPs (Figure 3C). The difference in proline abundance between UEPs and IEPs was significant in positions 4, 6 and 7 (Figure 3D). Of note, we also observed strong changes in the peptide anchor positions 2 and 9 that had not been apparent in the MIP vs MDP analysis. This can be explained by the relative overrepresentation of peptides presented by HLA-B among UEPs and by HLA-A in IEP, which is a consequence of the different levels of upregulation with IFNγ we described above. To control for this, we also separated the condition-exclusive peptide groups by their source HLA (Supplementary Figure 2C). Due to the lower peptide numbers there is more variation in the data, but the IFNγ-exclusive peptides showed a decrease in proline abundance in position 4 across different HLA allotypes.

Taken together, our approach to scrutinize the changes in the peptidome showed that IFNγ treatment results in an increased production of peptides with lysine, and a decreased production of peptides with small amino acids, at the N-terminus. Further to this, we found a decrease of peptides with proline in the center, mainly positions 4 and 5, a pattern that was present across all three PDOs and different HLAs, strongly suggesting that this effect is conserved across biological models.

We next sought to validate our results in an independent dataset. The immunopeptidomics and proteomics datasets from the breast cancer cell line MDA-MB-231(24) was appropriate for comparison as these are cancer cells that had also been treated with IFNγ for 48 hours. We applied the same MIP vs MDP analysis method to 9-mers (Figure 4A). This confirmed the underrepresentation of proline at positions 4-6 and 8 of MIPs (Figure 4B). Plotting the proline abundance between MIPs and MDPs, and statistical analysis with the Fischer’s exact test, showed statistically significant different in positions 3-6, and 8-9 (Figure 4C). We again observed a modest overrepresentation of lysine and arginine in position 1 of MIPs, however, this did not reach the Z-score threshold (Figure 4B).

**Figure 4.**
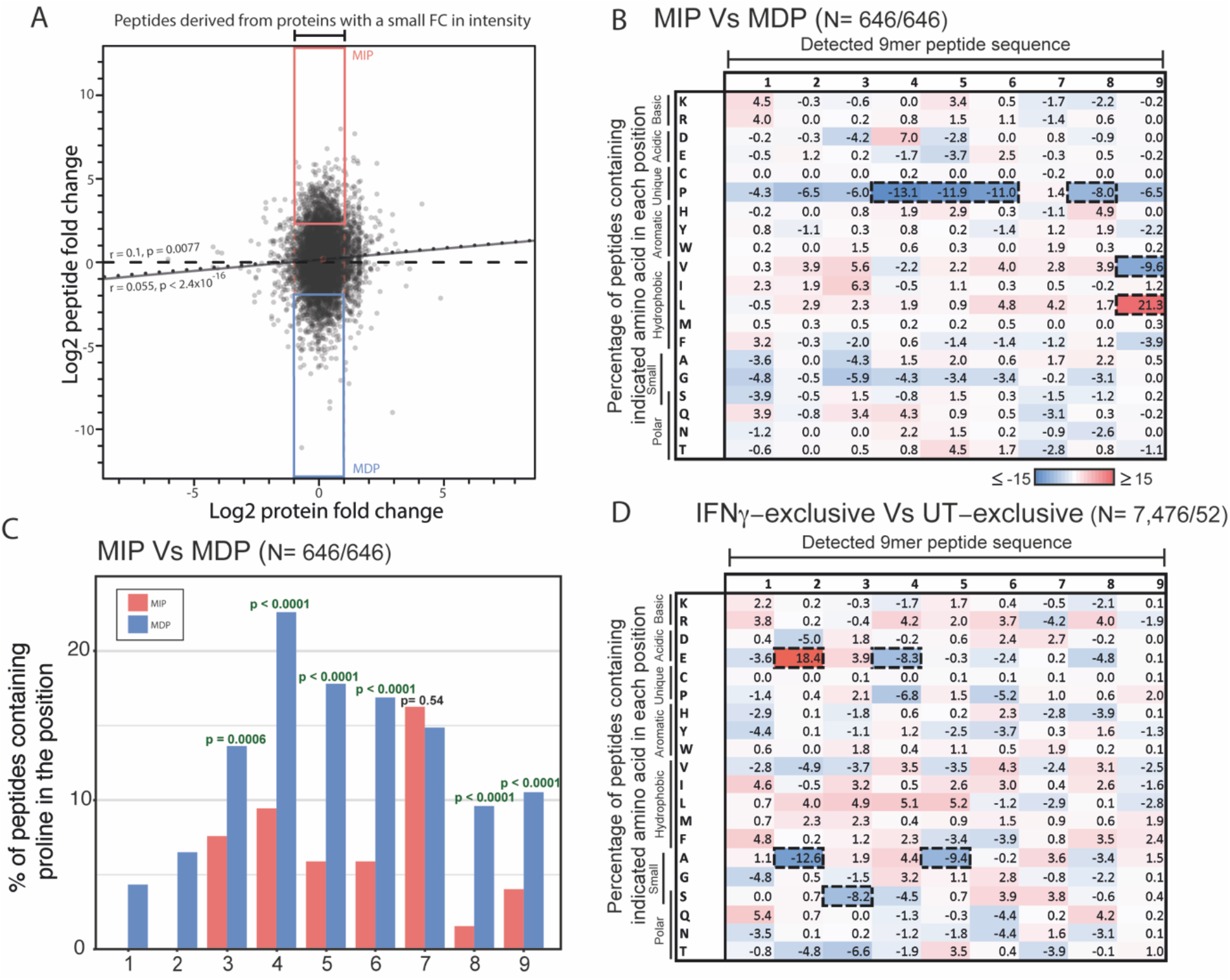
Validation of amino acid differences in the datasets from Goncalves et al. **A:** Correlation of proteomics fold-change between untreated and IFNγ conditions, against normalized immunopeptidomics fold-change. Regression line for all peptides marked by a solid black line, regression line for peptides derived from proteins with FC > 2 and < 0.5 marked with a dotted line. Spearman rank analysis used to investigate correlation. MIPs (top 10^th^ percentile peptide FC) and MDPs (bottom 10^th^ percentile peptide FC) derived from low fold-change proteins (0.5-2X fold-change) highlighted with red and blue boxes, respectively. **B:** Heatmap of the 9-mer peptide amino acid constituent changes between MIPs and MDPs, as highlighted in Figure A. Percentage of peptides with highlighted amino acid in each position were calculated for each group, then the percentage values for the MDPs were subtracted from the MIPs. **C:** A graph depicting the percentage of peptides with proline in each position for the MIPs and MDPs. **D**. Heatmap of the 9-mer peptide amino acid constituent changes between (IFNγ-exclusive peptides – IEPs) and (untreated-exclusive peptides – UEPs). Peptides were selected from ‘low fold-change’ proteins (0.5-2X fold-change). Percentage of peptides with highlighted amino acid in each position were calculated for each group, then the percentage values for the UEPs were subtracted from the IEPs. Z-score analysis was used for the heatmaps, and changes with a Z-score <-2.5 or >2.5 were highlighted with a thick dotted line. Fisher’s exact test was used for statistical analysis of the proline abundance values, with significant results are indicated in green.

When the UEP vs IEP analysis was applied, it showed a decrease in proline in positions 4 and 6 in IEPs (Figure 4D) which did not cross the Z-score threshold. This could be due to the small sample number of 50 UEPs compared to 7,476 IEPs. Overall, the validation of an underrepresentation of proline in MIPs strongly supports that peptides harboring proline residues in positions 4, 5 and possibly also 6-8, are specifically downregulated through IFNγ.

Proline has a unique impact on peptide secondary structures; its cyclic side chain gives the amino acid conformational rigidity, inducing a ‘kink’ of the amino acid sequence away from the proline residue which destabilizes secondary structures like helices and beta pleated sheets(37,38). We therefore investigated whether proline substitution impacts peptide affinity to or stability with its associated HLA.

We first assessed whether proline in position 4-8 influences the NetMHCpan4.0-predicted binding affinity to HLAs or the NetMHCstabpan1.0-predicted binding stability. Although binding affinity and stability are linked, they can differ, and stability may be more important for recognition by T-cell receptors(39). We sequentially replaced every amino acid in turn with proline, in a sample of 200 randomly selected 9-mer MS-detected peptides from our PDOs, to investigate the impact proline inclusion has on the NetMHCpan4.0 predicted affinity and NetMHCstabpan1.0 predicted stability (Figure 5 A-B). Replacing amino acids with proline was disadvantageous for peptide-HLA binding affinity and stability in the anchor residue positions P1-3 and P9. One exception was in the peptides which bind HLA-B35.08 in P2, where proline acts as an anchor residue, which saw increased affinity and stability. In contrast, substituting amino acids in P4-8 with proline had no strong effect on predicted binding affinity or stability, suggesting there is low specificity in these positions for peptide-HLA binding. To scrutinize this further, we selected all peptides containing a single proline and replaced it with either alanine or leucine (Figure 5 C-D and E-F). Alanine was selected as it eliminates side-chain interactions and doesn’t distort the confirmation of the main chain like proline. Leucine has similar properties to alanine, but it is a larger amino acid, so it is used where maintaining amino acid size may be important. Proline was infrequently detected in P1, 3, and 9 as it is less well tolerated in anchor positions for most HLA-I. The exception was again HLA-B35.08, which provides 94% of the detected peptides with proline in P2, due to its preference for proline as the anchor residue. Replacement of proline in P2 led to a large decrease in binding affinity and stability. Only small changes in peptide-HLA binding affinity and stability are seen in when replacing proline with alanine or leucine in P4-8; suggesting that proline in these positions doesn’t cause any notable structural changes, and doesn’t influence the affinity or stability of the peptides for their predicted HLA.

**Figure 5.**
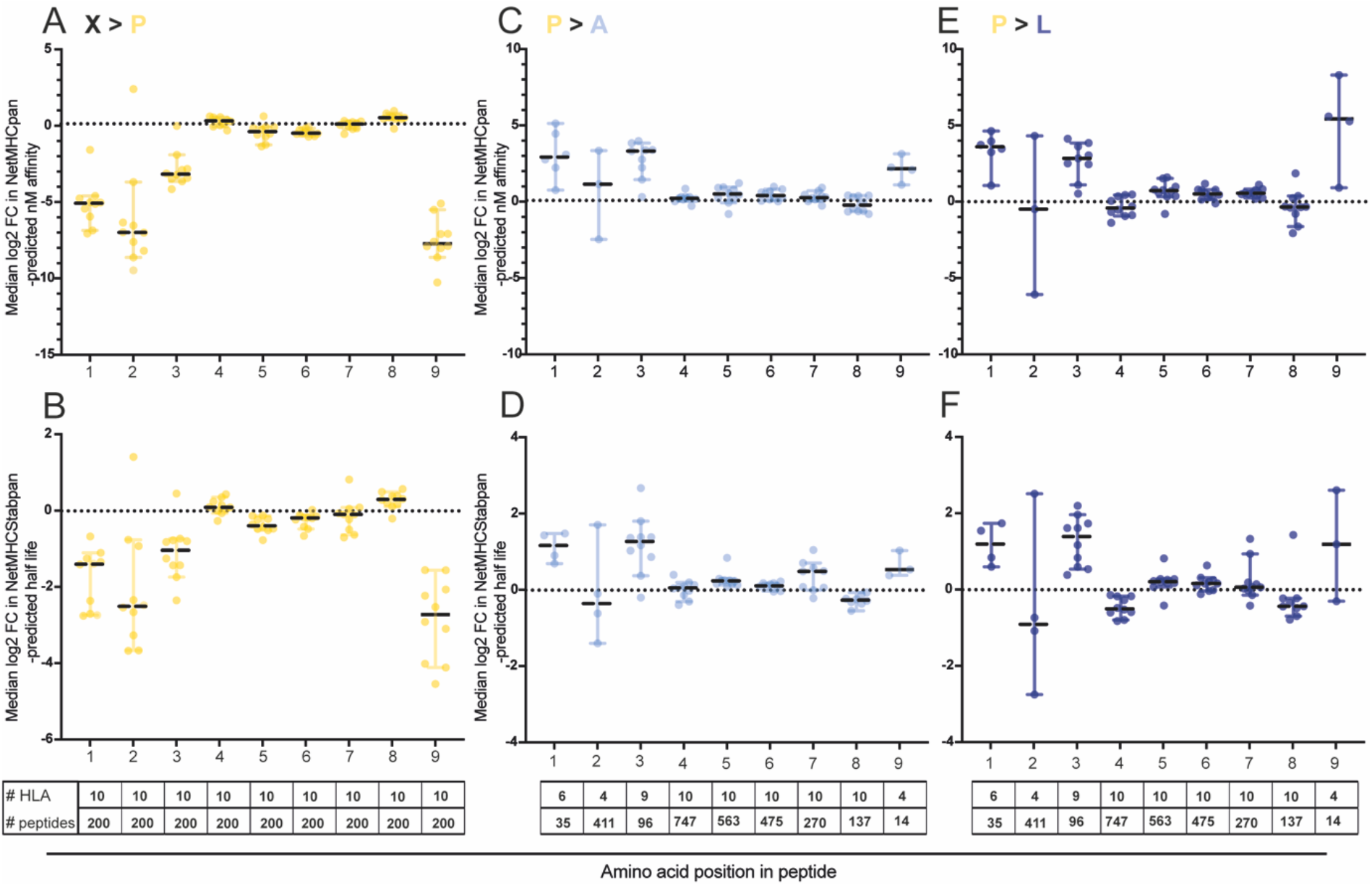
Simulating the impact of amino acid replacements on peptide affinity and binding stability to their cognate HLAs. **A:** Median log2 FC value from each HLA-A and -B from each PDO, demonstrating the impact of individually exchanging every single amino acid from detected peptides in positions 1-9 with proline, on NetMHCpan4.0-predicted nM affinity (N = 200 peptides per HLA). **B:** Median log2 FC value from each HLA-A and -B from each PDO, demonstrating the impact of individually exchanging every single amino acid from detected peptides in positions 1-9 with proline, on NetMHCStabpan1.0-predicted (N = 200 peptides per HLA). **C:** Median log2 FC value of each HLA-A and -B from each PDO, demonstrating the impact of exchanging proline, in detected peptides with a single proline, with alanine, on NetMHCpan4.0-predicted nM affinity (inidividual sample numbers annontated on the plots). **D:** Median log2 FC value of each HLA-A and -B from each PDO, demonstrating the impact of exchanging proline, in detected peptides with a single proline, with alanine, on on NetMHCStabpan1.0-predicted half life (inidividual sample numbers annontated on the plots). **E:** Median log2 FC value of each HLA-A and -B from each PDO, demonstrating the impact of exchanging proline, in detected peptides with a single proline, with leucine (inidividual sample numbers annontated on the plots), on NetMHCpan4.0-predicted nM affinity. **F:** Median log2 FC value of each HLA-A and -B from each PDO, demonstrating the impact of exchanging proline, in detected peptides with a single proline, with leucine (inidividual sample numbers annontated on the plots), on NetMHCStabpan1.0-predicted.

## Discussion

This study shows that peptide remodeling through IFNγ is complex and influenced by multiple distinct mechanisms. Up- or downregulation of proteins by IFNγ largely showed concordant abundance changes of corresponding peptides. However, an even larger fraction of peptides changed in cell surface abundance despite rather stable protein abundance. We and others have previously shown that this can be attributed, in part, to the increased chymotryptic activity of the immunoproteasome(21,23). Furthermore, IFNγ signaling disproportionally increased HLA-B compared to HLA-A surface expression which led to an increase in peptides presented on HLA-B. Moreover, demonstrating that even peptides originating from the same protein can show a mix of up- and downregulation, and that this is neither dependent on their HLA binding affinities, nor their location within the source protein, allowed us to isolate peptide-specific characteristics that affect their abundance in IFNγ conditions.

The most notable novel finding was the underrepresentation of proline in the core of MIPs and IEPs. It has previously been shown that proline in the core of the peptide sequence protects peptides from internal cleavage by endopeptidases and the proteasome(35,40). Known proline endopeptidases are DPP9, PREP, DPP8, DPP3, but these were not upregulated by IFNγ in our data. However, it is possible that the activity of such peptidases increases. An alternative theory is that the protective effect of proline is more relevant in the absence of IFNγ. This may be due to the lower abundance and activity of the antigen processing and presentation machinery, which may cause peptides to spend more time in the cytoplasm and ER before being shuttled to the cell surface, giving any single peptide more exposure to peptidases and hence, a higher probability of internal cleavage. Acceleration of peptide generation, processing, and loading onto HLA-I through IFNγ may allow peptides without proline to escape degradation. This may dilute proline-containing peptides within the peptide pool and explain their relative decrease or drop out.

Our insights into determinants of peptide abundance changes with IFNγ exposure could be useful to improve the design of cancer vaccines or TCR engineered therapies as it could enable the more accurate selection of peptides likely to be presented on patient tumors. For example, peptides presented on HLA-B, which are likely to increase in intensity when T-cells release IFNγ, may be preferable targets over those presented by HLA-A. Peptides with proline in positions 4 and 5, which favors peptide downregulation or even makes them undetectable in cancer cells exposed to IFNγ, can be avoided. Whether the overall increased in HLA surface expression or the remodeling of the immunopeptidome is more relevant for the critical role of IFNγ signaling for immunotherapy responses needs to be further investigated.

