## Supplementary materials for "Multifactorial remodeling of the cancer immunopeptidome by interferon gamma"

### Supplementary material for Newey et al.

#### Table of contents:

***Supplementary Methods:*** *Proteome analysis* **Page 2-4**

***Supplementary Table 1:*** *Summary statistics of most increasing and most decreasing peptides.* **Page 5**

***Supplementary Table 2:*** *Unique peptide count for untreated-exclusive peptides and IFN$\gamma$-exclusive peptides.* **Page 6**

***Supplementary Figure 1:*** *Effect of relative peptide position within protein on peptide abundance changes under IFN$\gamma$ treatment.* **Page 7-9**

***Supplementary Figure 2:*** *Additional internal validation of difference in amino acid features of peptides between untreated and IFN$\gamma$-treated conditions* **Page 10**

#### Supplementary Methods

**Proteome analysis**

Cell pellets were lysed with SDC lysis buffer [1% sodium deoxycholate, 100 mM triethylammonium bicarbonate (TEAB), 10% glycerol, 50mM NaCl] with Halt$^{TM}$ protease and phosphatase inhibitor cocktail (ThermoFisher Scientific). Cell pellet samples were completely homogenized with probe sonication (EpiShear) for 15 seconds at 40% power with 1 second on and 1 second off, heated at 90$^{o}$C for 5 minutes and then repeated the probe sonication. Proteins were quantified using Quick Start™ Bradford Protein Assay (Bio-Rad).

100 µg protein was taken from each sample and lysis buffer was added so each sample were at the same volume. Proteins were reduced with 10 mM tris(2-carboxyethyl)phosphine hydrochloride solution (TCEP, Sigma) at room temperature (RT) for 10 minutes and then alkylated with 5 mM iodoacetamide (IAA, Sigma) for 30 minutes at RT. Protein was then purified by 20% trichloroacetic acid precipitation. The pellet was resuspended in 100 mM TEAB buffer, and digested by 3.3 µg trypsin (Pierce, MS Grade) at a ratio of 1:30 (trypsin:protein by weight) at 37°C for 18 hours.

40 µg of protein digest were labelled with 0.5 mg TMTpro$^{TM}$ 16plex reagents (ThermoFisher Scientific) according to the manufacturer's instruction. After 1 hour incubation at RT and 15 min quenching by 4 µl of 5% hydroxylamine (ThermoFisher Scientific), the labelled samples were combined. Sodium deoxycholate was precipitated by adding formic acid (FA) (Honeywell Fluka). After centrifugation, the supernatant was collected and dried in Speedvac.

The sample were resuspended in 0.1% $NH_4OH$/100% $H_2O$, and fractionated on an XBridge BEH C18 column (2.1 mm i.d. x 150 mm, Waters) with an initial 5 min loading then linear gradient from 5% ACN/0.1% $NH_4OH$ (pH 10) – 35% $CH_3CN$ /0.1% $NH_4OH$ in 30 min, then to 80% $CH_3CN$ /0.1% $NH_4OH$ in 5 min and stayed for another 5 min. The flow rate was at 200 µl/min. Fractions were collected at every 42 seconds from retention time at 7.8 min to 50 min and then concatenated to 28 fractions and dried in SpeedVac. Samples were then resuspended in 0.5% FA for LC-MS/MS analysis.

**LC-MS/MS Analysis**

The LC-MS/MS analysis was on the Orbitrap Fusion Lumos mass spectrometer coupled with U3000 RSLCnano UHPLC system. All instrument and columns used below were from Thermo Fisher.

50% of peptides were injected. The peptides were first loaded to a PepMap C18 nanotrap (100 µm i.d. x 20 mm, 100 Å, 5 µm) at 10 µl/min with 0.1% FA/$H_2O$, and then separated on a PepMap C18 column (75 µm i.d. x 500 mm, 100 Å, 2 µm) at 300 nl/min with a linear gradient of 8-32% ACN/0.1% FA in 90 min /total cycle time at 120 min for each fraction. The data acquisition used standard data-dependant acquisition mode with a cycle time at 3 sec. The full MS scans (m/z 375-1500) were acquired in Orbitrap with a resolution at 120,000 at m/z 200, and the automatic gain control (AGC) was set at 400,000 with maximum injection time at 50 msec. The most abundant multiply charged ions (2+ ~ 5+) with intensity threshold at 5000 were isolated by quadrupole at the isolation window at 0.7 Da and then subjected to MS/MS fragmentation by Collision Induced Dissociation (CID) in ion trap at 35% normalized collision energy (NCE). The AGC was set at 10,000 and maximum injection time at 35 msec. The TMT report ions were detected by further fragmentation of the 5 most abundant fragment ions produced in MS2: they were isolated by synchronous precursor selection (SPS) method with the isolation width at 0.7 Da, and fragmented by higher energy collisionally activated dissociation (HCD) at 55% NCE, and detected in the Orbitrap in a scan range 100-500 m/z. The resolution was set at 50,000 at m/z 200, the AGC at 50,000 with maximum injection time at 86 msec. The dynamic exclusion was set 40 s with ± 10 ppm exclusion window.

**Mass Spectral Data Processing**

All raw files were processed in Proteome Discoverer 2.4 (Thermo Fisher) using the Sequest HT search engine to searched against reviewed Uniprot database of Homo Sapiens (Version February 2020) and contaminate database (from Thermo). Search parameters were: trypsin with 2 maximum miss-cleavage sites, mass tolerances at 10 ppm for the precursor, and 0.5

Da for the fragment ions; dynamic modifications of Carbamidomethyl (C), Deamidated (N, Q), TMTpro (K, peptide N-terminus) and Oxidation (M) and Acetyl (protein N-terminus). Search result was validated by Percolator with q-value set at 0.01 for the decoy database search, and only high confident PSMs (Peptide Spectrum Matches) were considered. Protein FDR was set at 0.01. Only master proteins were reported. For reporter ion intensity detection, the reporter ion quantifier node parameters were integration window tolerance 20ppm, integration most confident centroid for peak detection.  Only unique peptides were considered for quantification. TMTpro Quan value correction factor, provided by the manufacturer's certificate of analysis, was applied.  Co-isolation threshold was set at 100, reporter ions average S/N threshold at 3 and SPS mass matches threshold 55%. Report ions intensities were normalized by total peptide amount to correct the variation by for different protein loading in each channel, and then scaled on all average.

### Supplemental tables

Supplementary table 1. Statistics summary from our peptides which most increase and decrease in intensity, derived from proteins with a -1 to +1 log2 fold change, separated by NetMHCpan4.0-attributed HLA: unique peptide count, distribution and mean and median of immunopeptidomics FC.

| Category | PDO line | Source HLA | Peptide count | 1st Quartile | Median | Mean | 3rd Quartile |
|---|---|---|---|---|---|---|---|
| MIP | CRC-01 | HLA.A31.01 | 204 | 3.23 | 4.44 | 15.95 | 7.78 |
| MIP | CRC-01 | HLA.A32.01 | 137 | 5.34 | 7.73 | 17.50 | 13.09 |
| MIP | CRC-01 | HLA.B14.01 | 52 | 10.08 | 13.80 | 24.31 | 24.10 |
| MIP | CRC-01 | HLA.B27.05 | 164 | 10.60 | 14.50 | 23.14 | 24.43 |
| MIP | CRC-01 | HLA.C02.02 | 20 | 4.46 | 6.28 | 17.39 | 10.47 |
| MIP | CRC-01 | HLA.C08.02 | 56 | 4.07 | 5.72 | 7.67 | 8.27 |
| MIP | CRC-05 | HLA.A32.01 | 36 | 4.78 | 7.40 | 10.03 | 9.13 |
| MIP | CRC-05 | HLA.B40.01 | 92 | 7.14 | 8.89 | 19.21 | 12.60 |
| MIP | CRC-05 | HLA.C03.04 | 40 | 4.63 | 5.24 | 8.22 | 8.32 |
| MIP | CRC-04 | HLA.A03.01 | 188 | 4.07 | 5.45 | 20.05 | 8.70 |
| MIP | CRC-04 | HLA.A24.02 | 85 | 2.97 | 5.63 | 16.90 | 13.88 |
| MIP | CRC-04 | HLA.B18.01 | 31 | 5.20 | 6.56 | 24.15 | 15.30 |
| MIP | CRC-04 | HLA.B35.08 | 57 | 5.29 | 8.34 | 66.10 | 16.79 |
| MIP | CRC-04 | HLA.C04.01 | 5 | 23.14 | 26.29 | 46.77 | 34.53 |
| MIP | CRC-04 | HLA.C05.01 | 9 | 9.42 | 10.40 | 12.17 | 15.06 |
| MDP | CRC-01 | HLA.A31.01 | 204 | 0.08 | 0.11 | 0.11 | 0.14 |
| MDP | CRC-01 | HLA.A32.01 | 136 | 0.12 | 0.18 | 0.17 | 0.22 |
| MDP | CRC-01 | HLA.B14.01 | 52 | 0.17 | 0.25 | 0.23 | 0.30 |
| MDP | CRC-01 | HLA.B27.05 | 160 | 0.22 | 0.31 | 0.30 | 0.41 |
| MDP | CRC-01 | HLA.C02.02 | 19 | 0.11 | 0.16 | 0.17 | 0.23 |
| MDP | CRC-01 | HLA.C08.02 | 56 | 0.10 | 0.18 | 0.16 | 0.21 |
| MDP | CRC-05 | HLA.A32.01 | 36 | 0.12 | 0.17 | 0.16 | 0.21 |
| MDP | CRC-05 | HLA.B40.01 | 92 | 0.18 | 0.35 | 0.33 | 0.47 |

| | | | | | | | |
|---|---|---|---|---|---|---|---|
| MDP | CRC-05 | HLA.C03.04 | 40 | 0.08 | 0.15 | 0.14 | 0.20 |
| MDP | CRC-04 | HLA.A03.01 | 186 | 0.15 | 0.23 | 0.22 | 0.31 |
| MDP | CRC-04 | HLA.A24.02 | 84 | 0.09 | 0.15 | 0.13 | 0.18 |
| MDP | CRC-04 | HLA.B18.01 | 31 | 0.24 | 0.35 | 0.31 | 0.40 |
| MDP | CRC-04 | HLA.B35.08 | 57 | 0.21 | 0.31 | 0.27 | 0.35 |
| MDP | CRC-04 | HLA.C04.01 | 6 | 0.40 | 0.46 | 0.42 | 0.47 |
| MDP | CRC-04 | HLA.C05.01 | 9 | 0.50 | 0.53 | 0.49 | 0.54 |

Supplementary table 2. Unique peptide count for untreated-exclusive peptides and IFNγ-exclusive peptides from our 3 CRC PDOs, separated by NetMHCpan4.0-attributed HLA.

| | **Untreated-exclusive peptides** | **IFNγ-exclusive peptides** |
|---|---|---|
| CRC-01 HLA.A31.01 | 162 | 170 |
| CRC-01 HLA.A32.01 | 88 | 148 |
| CRC-01 HLA.B14.01 | 15 | 71 |
| CRC-01 HLA.B27.05 | 44 | 345 |
| CRC-01 HLA.C02.02 | 24 | 33 |
| CRC-01 HLA.C08.02 | 89 | 109 |
| CRC-05 HLA.A32.01 | 12 | 14 |
| CRC-05 HLA.B40.01 | 48 | 163 |
| CRC-05 HLA.C03.04 | 17 | 44 |
| CRC-04 HLA.A03.01 | 200 | 273 |
| CRC-04 HLA.A24.02 | 156 | 115 |
| CRC-04 HLA.B18.01 | 66 | 84 |
| CRC-04 HLA.B35.08 | 142 | 115 |
| CRC-04 HLA.C04.01 | 5 | 43 |
| CRC-04 HLA.C05.01 | 26 | 32 |

Supplementary Figure 1

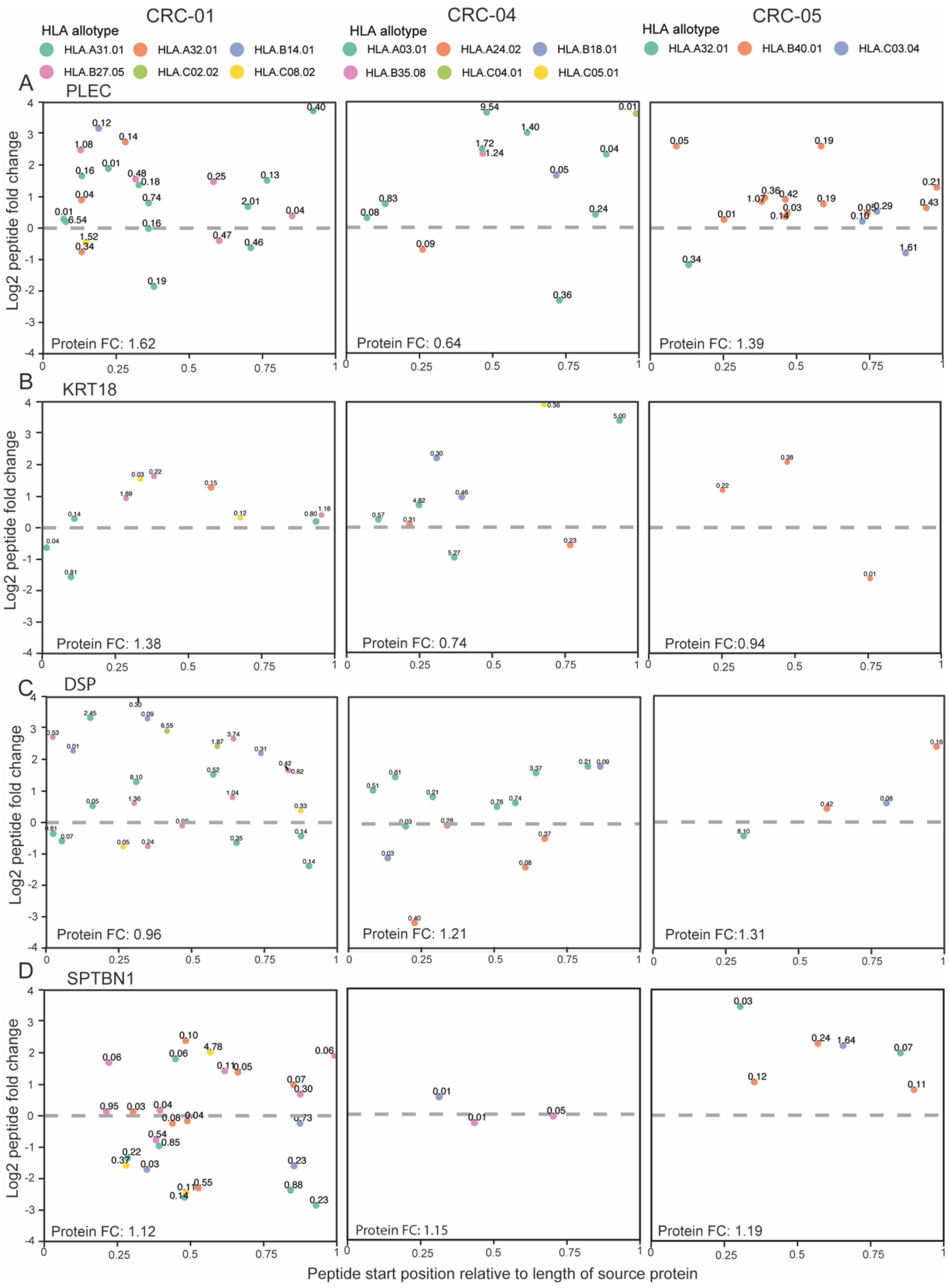

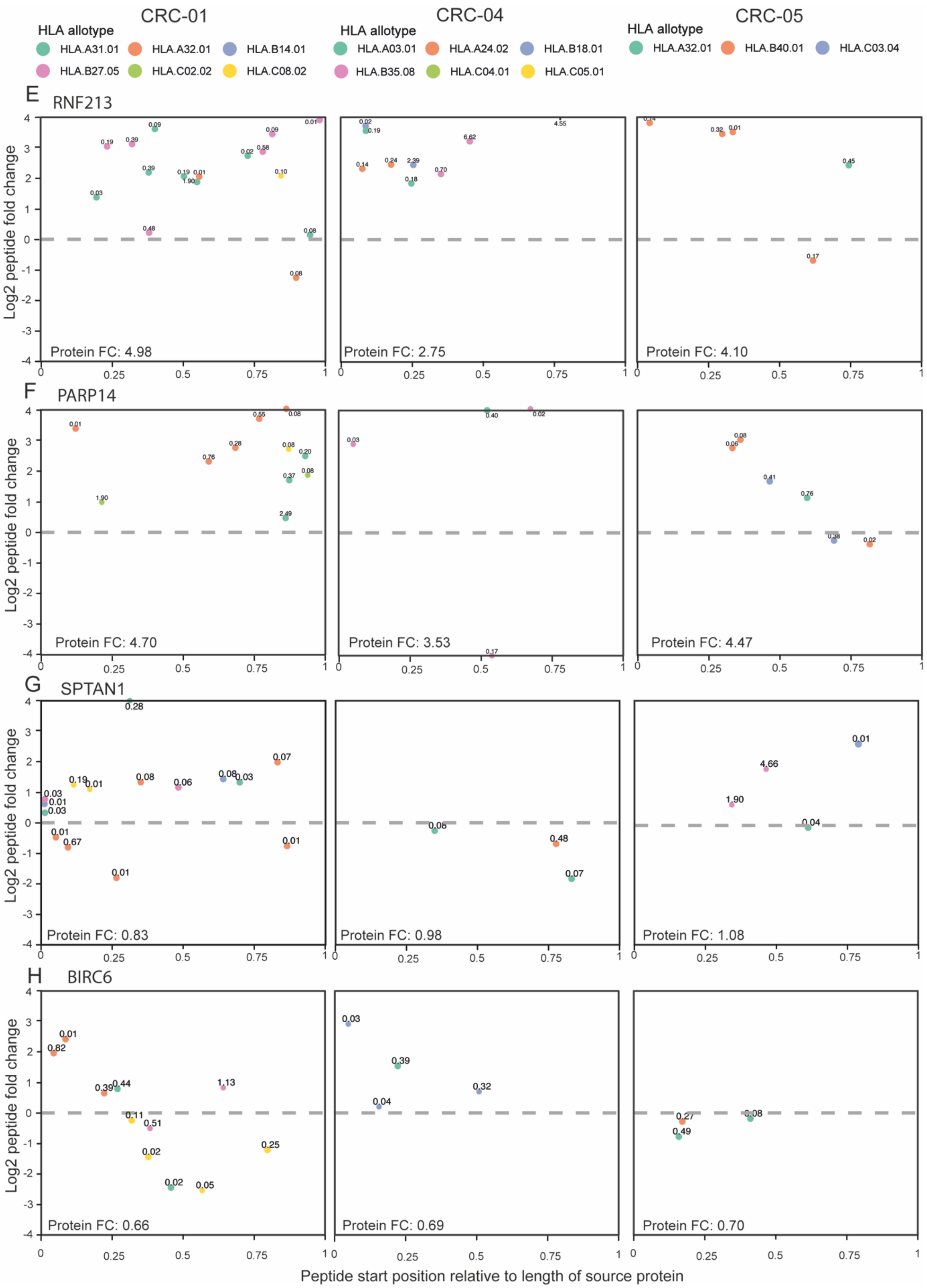

CRC-01
CRC-04
CRC-05
HLA allotype
HLA.A31.01 HLA.A32.01 HLA.B14.01 HLA.B27.05 HLA.C02.02 HLA.C08.02
HLA.A03.01 HLA.A24.02 HLA.B18.01 HLA.B35.08 HLA.C04.01 HLA.C05.01
HLA.A32.01 HLA.B40.01 HLA.C03.04
E RNF213
Protein FC: 4.98
Protein FC: 2.75
Protein FC: 4.10
F PARP14
Protein FC: 4.70
Protein FC: 3.53
Protein FC: 4.47
G SPTAN1
Protein FC: 0.83
Protein FC: 0.98
Protein FC: 1.08
H BIRC6
Protein FC: 0.66
Protein FC: 0.69
Protein FC: 0.70
Log2 peptide fold change
Peptide start position relative to length of source protein

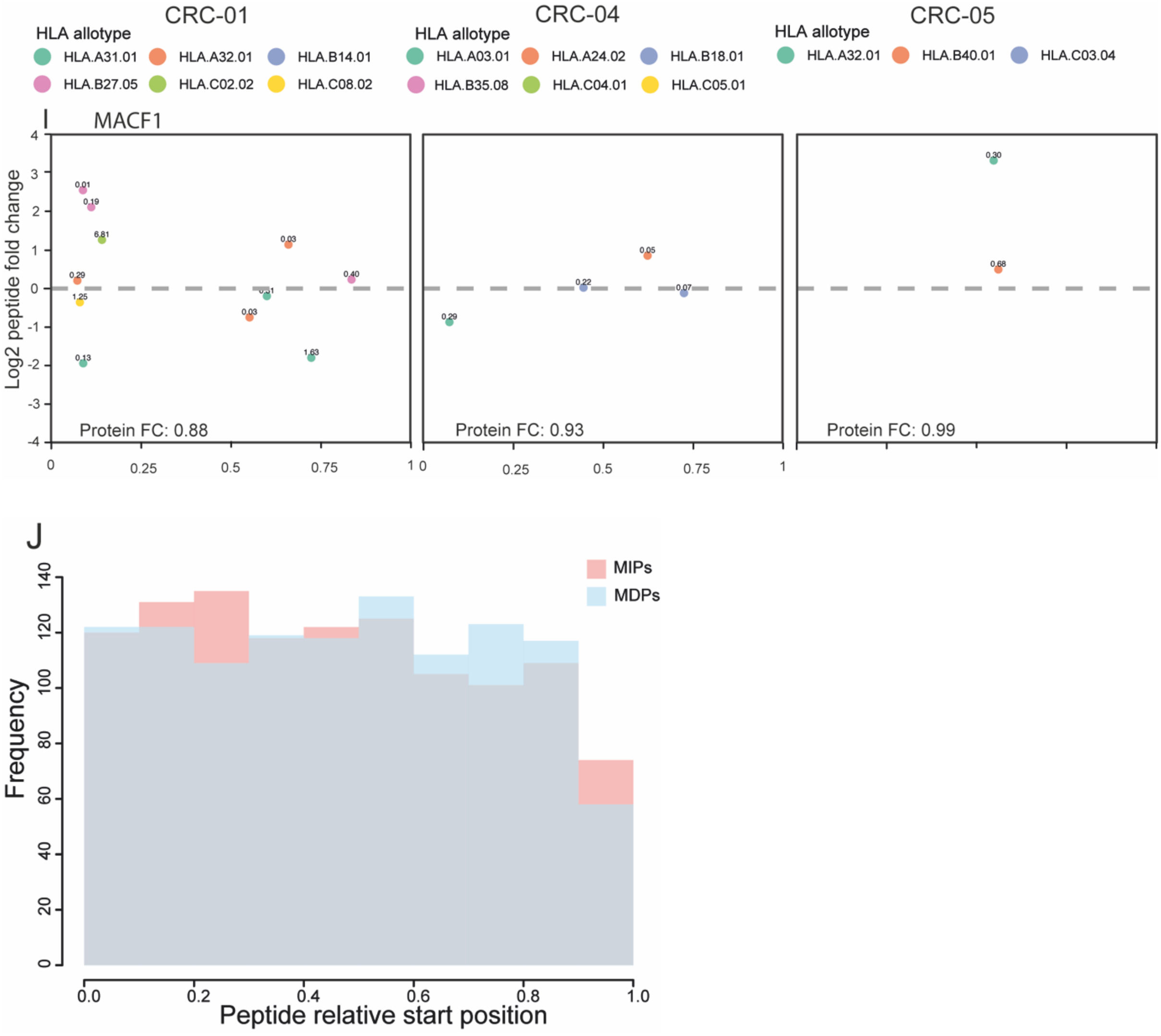


**Supplemental figure 1*. Effect of relative peptide position within protein on peptide abundance changes under IFNγ treatment.*** Log2 change in peptide intensity between untreated and IFNγ conditions for peptides derived from the single source protein, plotted against the relative position of the peptide in protein. Long proteins with the most peptides across the 3 PDOs selected. Points color-coded by their NetMHCpan4.0-predicted source HLA, with the NetMHCpan4.0 predicted rank annotated above. Protein fold change for the source protein in each organoid noted at the bottom of each plot. **A-I:** PLEC, KRT18, DSP, SPTBN1, RNF213, PARP14, SPTAN1, BIRC6, MACF1. **J:** Frequency distribution of peptide relative distribution from all MIPs against all MDPs.

#### Supplementary Figure 2

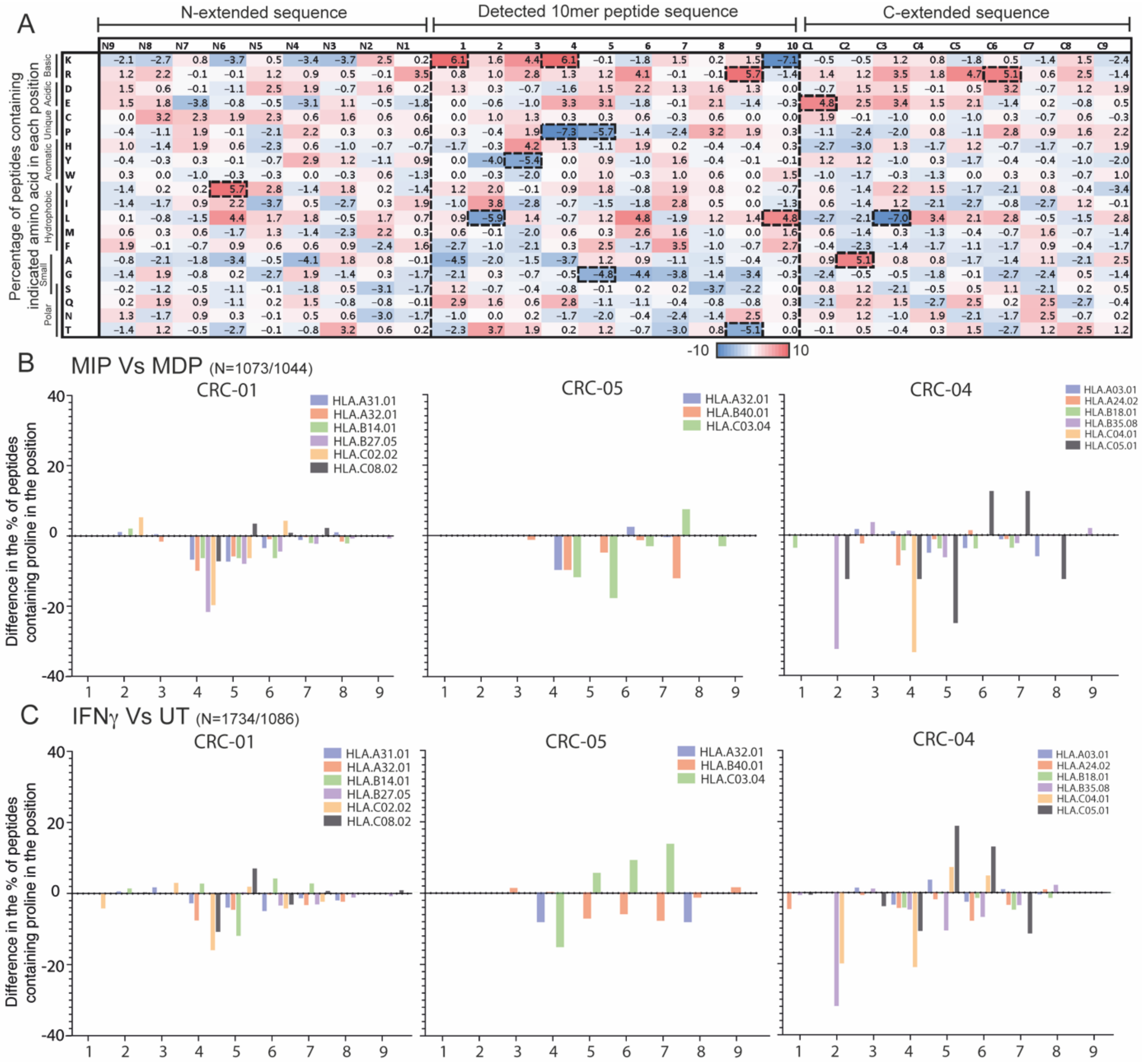


**Supplemental figure 2.** ***Additional internal validation of difference in amino acid features of peptides between untreated and IFN$\gamma$-treated conditions.*** **A:** Heatmap of the 10-mer peptide amino acid constituent changes between MIPs and MDPs. Peptides were selected from ‘low fold-change’ proteins (0.5-2X fold-change). Percentage of peptides with highlighted amino acid in each position were calculated for each group, then the percentage values for the MDPs (N=333) were subtracted from the MIPs (N=330). **B:** A graph depicting the change in the percentage of peptides with proline in each position between MIPs and MDPs, split by PDO and by NetMHCpan4.0-predicted source HLA (peptide numbers detailed in Supplementary Table 1 and 2). **C:** A graph depicting the change in the percentage of peptides with proline in each position between IFNγ-exclusive peptides and untreated-exclusive peptides, split by

PDO and by NetMHCpan4.0-predicted source HLA (peptide numbers detailed in Supplementary Table 3).
